# Modulation of *rpoS* fitness by loss of *cpdA* activity during stationary-phase in *Escherichia coli*

**DOI:** 10.1101/460451

**Authors:** Savita Chib, Aswin SaiNarain Seshasayee

## Abstract

Experimental evolution of *Escherichia coli* in one month long stationary-phase in lysogeny broth batch cultures repeatedly selected mutations in the genes for the stationary-phase sigma factor RpoS and the cAMP phosphodiesterase CpdA. The founder strain carried a previously identified allele of *rpoS*, referred to as *rpoS819,* a partially functional variant that confers growth advantage in stationary-phase (GASP). The 46 base duplication at the 3’ end of *rpoS819* produces a longer protein present at very low levels compared to wild type RpoS. A new *rpoS* variant *rpoS92*, carrying a re-duplication of the original duplication in *rpoS819,* arose during the first week of our evolution experiment. In *rpoS92*, an in-frame stop codon truncated RpoS819 creating a shorter RpoS92 whose levels are restored to that of wild type RpoS. Transcription profiling of *rpoS92* indicated a shift in gene-expression to that of wild-type *rpoS*, reversing some of the expression trends of *rpoS819*. Δ3*cpdA*, carrying an in-frame three base deletion, had arisen late in our evolution experiment. It is a loss of function mutation, which elevates cAMP levels. Using mixed culture competition experiments, we demonstrate that *rpoS92* confers GASP, whereas Δ3*cpdA* confers relatively modest GASP in comparison to the ancestral *rpoS819*. Δ3*cpdA* mediates epistatic repression of *rpoS92* GASP. The original survivor carrying both rpoS92 and Δ3*cpdA* besides other mutations displays robust GASP, highlighting the role of these additional mutations in reversing the epistatic interaction between Δ3*cpdA* and *rpoS92.* In 10- and 20-day old spent media, there is a reduction in the competitive fitness of *rpoS92,* which is arrested by Δ3*cpdA.* Thus the activity of RpoS fluctuates via genetic mutations in deep stationary phase, and additional mutations in CpdA helps modulate the competitive fitness of RpoS variants.

## Introduction

Bacterial growth in nature is often described as a feast and famine lifestyle (1). Mammalian gut - the primary habitat of *Escherichia coli* - is a mixed substrate niche, whereas extra-intestinal secondary habitats are typically resource limiting (2–5). Additionally, constant exposure to environmental insults such as changes in pH, temperature, salinity and intense competition for nutrients and other resources exact a balance between growth and survival (6–8). Savageau predicts that the average doubling time of *E. coli* in the natural environment is 40 hours, which is in stark contrast to the 20-30 minutes routinely observed in the laboratory during exponential growth under rich media conditions (9).

Micro-organisms including *E. coli* populations can survive long periods of nutrient limitation. *E. coli* survival during prolonged stationary-phase is illustrated by the phenomenon of <u>G</u>rowth <u>A</u>dvantage in <u>S</u>tationary-<u>P</u>hase (GASP). During GASP, genetic variants called GASP mutants arise and proliferate in the primarily non-growth environment of prolonged stationary-phase, presumably by extracting nutrients from dead cell debris (10–15).

One of the first GASP mutations identified from a starving *E. coli* population was a mutation in the stationary-phase sigma factor gene *rpoS* (10). RpoS (also called SigmaS or Sigma38) drives transcription of more than 15% of genes, many of which have roles to play in the general stress response, directly or indirectly (16–19). Despite its central role in general stress response, *rpoS* variants with attenuated activity have been observed frequently during prolonged phases of slow growth (8, 10, 22). Strains with attenuated *rpoS* display an improved ability to utilize poor carbon sources such as succinate and acetate but with a suboptimal stress response, whereas the opposite is true for strains with abundant RpoS (6, 8, 11, 21, 23-25). Thus, polymorphism at *rpoS* represent trade off between self-preservation and nutritional competence (6, 8, 26).

Nutritional competence on alternative carbon source is determined by the small signaling molecule cyclic Adenosine MonoPhosphate (cAMP), which is synthesized by the adenylate cyclase CyaA and hydrolyzed by the cAMP phosphodiesterase CpdA (27-29). cAMP forms a complex with the transcription factor Cyclic AMP Receptor Protein (CRP) (30). The active cAMP-CRP complex directly controls the expression of metabolic genes that are required for the utilization of alternative carbon resources (28, 31-33). For example, *cyaA* and *crp* mutants are compromised in their ability to use secondary sugars and amino acids as sole carbon source (34). cAMP-CRP has been implicated in both positive and negative regulation of RpoS (34-37). A large fraction of RpoS core promoters have cAMP-CRP binding site/s and nearly half of these sites are located such that they are likely to place the transcription factor in a repressive position (28, 38). Overlap between the RpoS and the cAMP-CRP regulons, and their role in general stress response and alternative carbon resource metabolism may determine the survival and proliferation of bacteria in resource deficient environments.

Previously, we reported the mutation spectrum of *E. coli* K12 populations maintained in long-term stationary phase (LTSP) after growth in lysogeny broth (39). One of the key observations from our study was strong genetic parallelism at *rpoS* and *cpdA* loci. Here, we seek to determine the role of *rpoS* and *cpdA* mutations in growth and survival during prolonged stationary-phase by: i) functional characterization of the mutant alleles; ii) mutation reconstruction and competition experiments to assess the fitness in specific genetic and environmental contexts; and (iii) gene-expression changes towards understanding the molecular basis of GASP.

## Methods

### Bacterial Media and Growth Conditions

Various *E. coli* strains were cultured in Lysogeny broth (LB) at 37 °C at 200 rpm. For determining viable counts, cultures were plated on LB agar medium supplemented with 35 μg of kanamycin (Kan)/ml, 100 μg of ampicillin (Amp)/ml, 10 μg of tetracycline (Tet)/ml, or 15 μg of chloramphenicol (Cam)/ml when required. Competition experiments were performed as described below.

### P1 transduction

P1 phage transduction were performed by standard protocol described in Miller (40).

### Competition Assays

For a standard stationary-phase mixed culture competition experiment, monocultures of the two competing strains were grown in 5 ml of LB at 37 °C with aeration for 12 hours. These overnight grown cultures were mixed in the desired proportion reciprocally (1:1000) and incubated under the same conditions without the addition of fresh medium. The two populations marked with two different antibiotic resistance markers were tracked by plating on LB agar plates containing the respective antibiotics every-day of the competition experiment duration. For competitions in conditioned medium, the parent strain ZK819 was grown for 21 days at 37 °C with periodic addition of sterile-distilled water to compensate for evaporation. Twenty-one day old culture was pelleted and supernatant was filtered twice through a filter with pore size 0.2 micron (Sartorius) to remove cells. To replicate the competitions in 21 day old conditioned medium, the overnight grown monocultures of competing strains were pelleted at 4000 rpm for 10 minutes. The supernatant was discarded and the cell pellets were resuspended in the conditioned medium. Cultures were mixed in 1:1000 proportion reciprocally and the competing populations were monitored as described above.

### Statistical Data Analysis

All the data were analyzed using the statistical programming language R (v3.4.4). Paired t-test between the slopes of the four replicates of two competing populations was performed to test the significance of the differences and P-values for each comparison is reported. To compare the slopes of a population across two competing sets, standard t-test was done. P < 0.05 was considered significant.

### Construction of cpdA deletion strain

Gene knockouts were generated by the procedure of Datsenko and Wanner (41). Hybrid primers with 36-bp extensions homologous to the N terminus and C terminus of the *cpdA* locus spanning positions +2 and +445 were designed for the PCR amplification of kanamycin resistant gene flanked by FLP recognition target sites. F*cpdA-kan* (5′- TATTAGCGTCGTGAAACCTAAGGACACCATTTGGAAGTAGGCTGGAGCTGCTTCG-3′) and R*cpdA*-*kan* (5′- TGAAACCGTGTAAATAAAGAAGCGTAGACATCAGTACATATGAATATCCTCCTTA-3′) encoded by plasmid pKD4. The PCR amplicon was used to replace the *cpdA* locus with the selectable Kan resistance gene (*kan*) by homologous recombination aided by phage lambda recombinase expressed from the helper plasmid pKD46. The deletion was confirmed by PCR using Forward primer FCpdA 5’- CAGAACGCCATACGTTGCTGCTGCTGC-3’ and reverse primer RCpdA 5’- GTATCAGGTTGGAAACGTGTGT-3’. Whenever necessary, the Kan resistance cassette was eliminated using the helper plasmid pCP20 expressing FLP recombinase, which leaves a scar of 80 nucleotides at the deletion site.

### Western Blotting

For detection of RpoS levels in-vivo, cells from desired bacterial culture grown in LB at 37 °C to an optical density at 600 nm (OD_600_) of 1.0 were collected and washed in Phosphate Buffer-Saline. Total protein was obtained by sonication and quantified using bicinchoninic acid assay (BCA Protein Assay Kit, Thermo Scientific). Total cellular proteins were electrophoresed using SDS–15% PAGE for three hours at 120 V and transferred onto a nitrocellulose membrane. *E. coli* anti-RpoS monoclonal antibody (Neoclone) was used to probe RpoS and the signal was detected using horseradish peroxidase (HRP)-conjugated anti-mouse IgG (Sigma). HRP luminescence was detected using West Dura reagent (Thermo Scientific). Digital images of the blots were obtained using an LAS-3000 Fuji imager. The level of GroEL, an internal loading control, was detected by using Rabbit polyclonal anti-GroEL antibodies (abcam).

### Cyclic adenosine monophosphate (cAMP) estimation

Estimation of intracellular cAMP levels was carried out using the cyclic AMP Select EIA kit (501040; Cayman Chemical). Cells grown till stationary phase (~12–15 hr in LB) were harvested by centrifugation at 13,000 × *g* for 1 min. Cells were immediately transferred onto ice to prevent breakdown of cAMP by phosphodiesterases. Cell pellets were washed once with TBST (20 mM Tris, 150 mM NaCl, 0.05% Tween 20, pH 7.5), resuspended in 0.05 N HCl and boiled for 5 min to extract cAMP. Cells were then spun down at 14,000 × g and the supernatant containing cAMP was collected. Estimation of cAMP was carried out according to the kit’s instructions with the exception that the provided cAMP standard was diluted in 0.05 N HCl to generate the standard curve, since HCl was used for the extraction process.

### DNA and RNA Sequencing

Genomic DNA was isolated from the original survivor Sur_Δ3*cpdA* clone using GenElute bacterial genomic DNA kit (NA2120; Sigma-Aldrich) according to the manufacturer’s protocol. The quality of genomic DNA was assessed using Nanodrop UV absorbance ratios as well as on 0.8% agarose gel. DNA was quantified using the Qubit double-stranded DNA high-sensitivity assay. Paired end whole genome sequencing library was prepared using Illumina’s TruSeq Nano DNA LT library preparation kit (FC-121-4001; Illumina) and run on Illumina’s MySeq platform with the sequencing depth of 2X150. The reads were mapped to the reference *E. coli* genome W3110 (GenBank accession no. NC_007779.1) and mutations were called using the breseq pipeline, which uses Bowtie for sequence alignment (42, 43).

For RNA Sequencing, two biological replicates for each strains were grown in LB at 37 °C at 200 rpm. Samples were harvested at 16 hours time point and immediately processed for total RNA isolation using Trizol method (15596018; Invitrogen). Dnase treated RNA samples were enriched for mRNA by depleting ribosomal RNA using the Ambion Microbe Express Kit (AM1905; Invitrogen). Illumina TruSeq RNA library prep Kit v2 was used to prepare single end sequencing libraries which were run on Illumina HiSeq 1000 platform. Each sample is sequenced to 50X read depth. RNA-Seq reads were aligned to W3110 genome (GenBank accession no. NC_007779.1) using SAMtools which uses BWA for sequence alignment (44, 45). Readcounts per gene were generated from SAM files using a custom perl script. Subsequently the R package DESeq was used to call differentially expression of genes (46). All subsequent data analyses on this set of mutations were performed using the statistical programming language R (v3.4.4).

## Results

### Genetic parallelism at rpoS and cpdA during LTSP evolution

In a previous study, we had sequenced genomic DNA from multiple long term stationary phase (LTSP) populations of *E. coli* and found evidence of genetic parallelism in which the same gene acquired mutations in multiple independent lines. Among such genes were *rpoS* and *cpdA* (39). RpoS is a general stress response sigma factor and a frequent target of mutation and selection under complex environments involving prolonged periods of nutrient limitation (6, 8, 10). The founder *E. coli* strain ZK819 used in our evolution experiment carries an *rpoS* allele, referred to as *rpoS819,* which shows attenuated activity. *rpoS819* was naturally selected during prolonged starvation in lysogeny broth batch culture and was shown to confer growth advantage in stationary-phase (10). We observed that all five independent replicate populations had diversified at the *rpoS* locus giving rise to four *rpoS* populations (39). These were (a) *rpoS819*; (b) an A-to-G substitution 21 bp upstream of *rpoS* in all samples including the parent; (c) *rpoS* with a 92 base duplication (*rpoS92*); and (d) *rpoS* with the original 46 base duplication as well as a 92 base duplication separated by 4 bases.

The second locus carrying mutations across multiple replicates is *cpdA* which encodes a phosphodiesterase that breaks down cAMP (29). Four out of five replicate populations had accumulated a variety of mutations in *cpdA*. Six cpdA genetic variants were observed across multiple populations: (a) *cpdA*:Δ3:bp486-488; (b) CpdA:L147P; (c) *cpdA*Δ3:bp116-118, (d) CpdA:S222L; (e) *cpdA*:Δ1:bp 356; and (f) *cpdA*:+CTC bp 622.

We isolated *rpoS92* and two independent *cpdA* clones - *cpdA*:Δ3:bp486-488 and *cpdA*:L147P - by screening frozen stocks of the evolved population saved during our previous LTSP experiment. We verified the genotype status of *rpoS* and *cpdA* using PCR amplification and Sanger sequencing. From whole genome re-sequencing data for these purified clones, we noted the presence of the *rpoS92* allele in both the *cpdA* clones along with additional mutations. To investigate the fitness potential of *cpdA* mutations, we selected the *cpdA*:Δ3:bp486-488 variant referred to as Δ3*cpdA* from an original survivor Sur_Δ3*cpdA.* Sur_Δ3*cpdA* was isolated from a Day 21 sample of the LTSP line referred to as ZK819.3 in our previous work (39). The list of mutations found in Sur_Δ3*cpdA* genome by whole genome short-read sequencing are given in Supplementary Table 1

### rpoS92 confers GASP in stationary-phase

The *rpoS92* allele arose early in our evolution experiment and steadily increased to nearly 40% (39). It had appeared in all the replicate populations by the end of the first quarter of our one month long experiment. Simultaneously, the ancestral *rpoS819* allele frequency declined in all replicates except one wherein it roughly maintained its starting frequency (39). To determine the GASP phenotype of *rpoS92*, we transduced the *rpoS92* allele to the ancestor ZK819 to create ZK92. To set up a stationary-phase competition, overnight grown monocultures of ZK92 and the ancestor ZK819 were mixed in 1:1000 ratio reciprocally. Growth trends of the strains in mixed culture over four days of competition showed that ZK92 minority counts increased by ~11 fold whereas the ancestor ZK819 majority declined by ~31 fold. This closes the gap in size between the populations, indicating that *rpoS92* confers growth advantage in stationary-phase (Fig. 1A; paired t-test comparing the slopes of the competing populations, N = 4, P = 0.003). In the reciprocal mix, the ZK92 majority decline was ~10 fold whereas the ancestor ZK819 minority counts dropped by ~50 fold (Fig. 1B; paired t-test comparing the slopes of the competing population, N = 4, P = 0.04).

**Figure 1:**
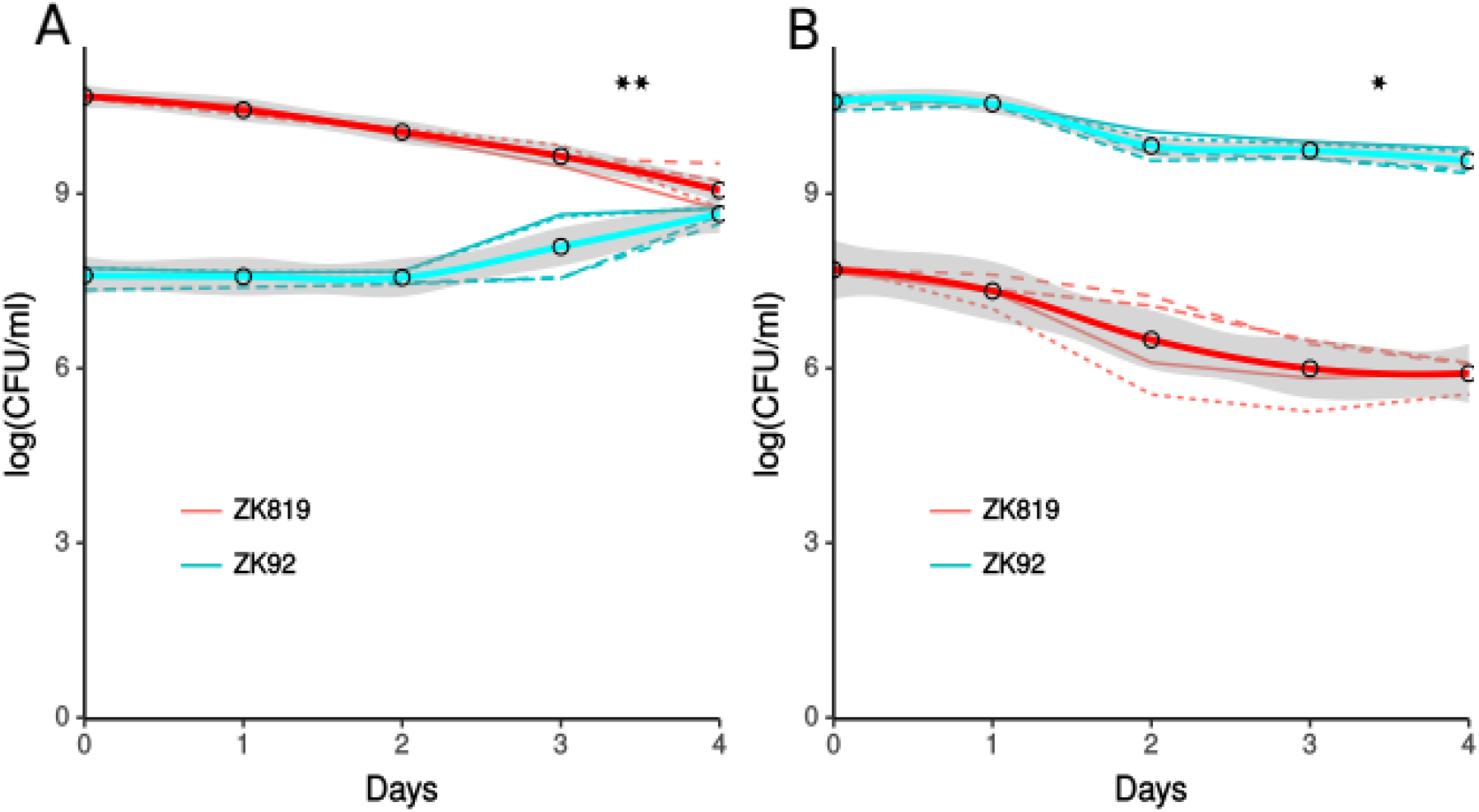
Stationary-phase competition between ZK92 (blue) and ZK819 (red) (A and B). One-day-old cultures of the two strains were mixed in ratios of 1:1,000 (A) and 1,000:1 (B). The mixed cultures were maintained in the original LB medium without the addition of fresh medium, and the growth of the two populations was monitored by scoring the number of CFUml^−1^. **P<0.005,*P<0.05.The thick curve is the LOE line fitted to the data.

**Table 1.**

| Strain | Genotype | Source |
| --- | --- | --- |
| ZK819 / Ancestor | ZK126 (W3110 $\Delta lacU169$) *rpoS819Sm<sup>r</sup>* | (10) |
| ZK92 | ZK819 *rpoS92 tna::Tn10Tet<sup>r</sup>* | This study |
| ZK$\Delta 3cpdA$ | ZK819 *cpdA*:$\Delta 3$:bp486-488 *mda*::Kan<sup>r</sup> | This study |
| ZK92$\Delta 3cpdA$ | ZK819 *rpoS92 cpdA*:$\Delta 3$:bp486-488 *mda*::Kan<sup>r</sup> | This study |
| ZK819*cpdA*<sup>nul</sup> | ZK819 *cpdA*::Kan<sup>r</sup> | This study |
| Sur_$\Delta 3cpdA$ | Original Survivor / ZK819 *rpoS92 cpdA*:$\Delta 3$:bp486-488 (and additional mutations) | This study |
| Sur_$\Delta 3cpdA$<sup>wt</sup> | Original Survivor *cpdA*<sup>wt</sup> | This study |

### RpoS92 is a truncated version of RpoS819

The ancestral *rpoS* allele *rpoS819* has a 46 base duplication at the 3’ end causing a shift in the open reading frame of *rpoS*. This 46 base duplication results in a longer RpoS variant in which the last four amino acids of wild-type RpoS are replaced with a stretch of 39 amino acids (Fig.2A & B). Sequence analysis of the new allele *rpoS92* revealed the reduplication of the original 46 base duplication (Fig. 2A). In *rpoS92*, the second duplication introduces an early stop codon that truncates RpoS819 (Fig. 2A & B). We probed the cellular levels of the RpoS polypeptide variants using anti-RpoS antibodies (Fig. 2C). RpoS819 indeed is the longest polypeptide, but present at very low levels (Fig. 2C, Lane 3). RpoS92 is noticeably shorter than RpoS819 and shows higher expression than RpoS819, to a level comparable to (or even higher than) that of wild-type RpoS (Fig. 2B, Lane 4). Differential gene-expression analysis of RpoS targets in *rpoS92* and *rpoS819* allelic background using RNA-seq shows that *rpoS92* has partially restored the *rpoS* regulon expression to that of wild-type *rpoS* (Fig. 2D). Thus the activity of RpoS fluctuates – through genotype alterations - in deep stationary phase.

**Figure 2:**
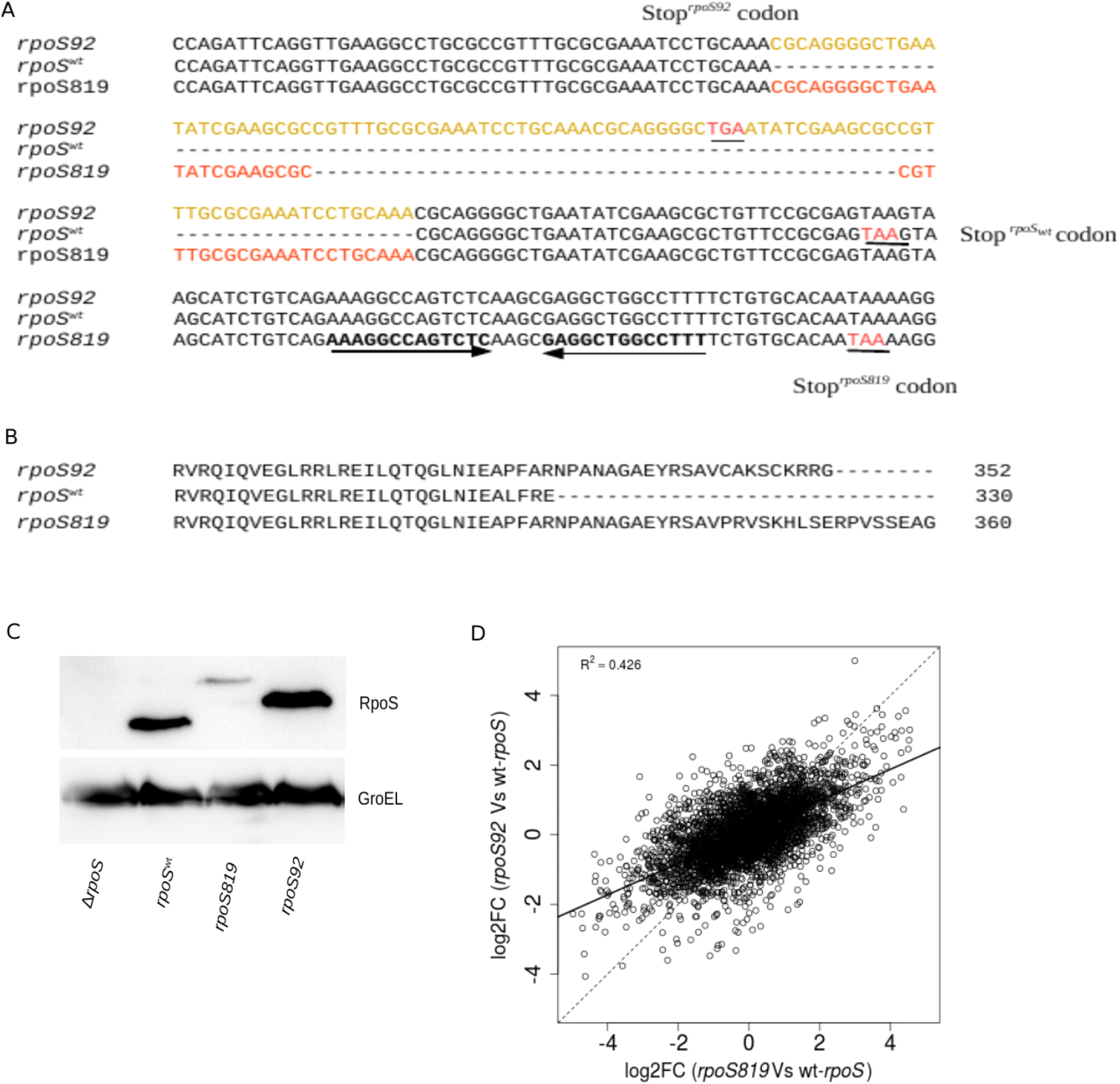
(A) Nucleotide sequence of the 3′ end of wild-type rpoS, rpoS819, and rpos92. The 46 base duplication in rpoS819 results in a frame-shift. The next stop codon is highlighted in read and preceding to inverted repeats-underlined by inverted arrows. (B) Putative sequence of RpoS819, RpoS92 polypeptides with extended C-terminal. Immunoblot analysis of RpoS polypeptides variants. (C) Total cell extracts of the strains ZK126 rpoS null (lane 1), ZK126 (lane 2) and ZK819 (lane 3) and ZK92 (lane 4) were probed using polyclonal anti-RpoS antibody. Anti-GroEL antibody was used to detect the loading control GroEL. (D) The scater plot representing the log2Fold Changes of gene-expression in rpoS92 andrpoS819 compared with wild-type rpoS gene-expression stationary-phase.

### The original cpdA survivor Sur_Δ3cpdA displays GASP

We tested the fitness of original Sur_Δ3*cpdA* against the ancestor ZK819 in mixed culture competition assay. By the end of three days of competition, Sur_Δ3*cpdA* minority counts had increased by ~32 folds while the ancestor majority counts dropped by ~806 folds (Fig. 3A; paired t-test comparing slopes of competing populations, N = 4, P = 2.21 x 10^−06^). This establishes the GASP phenotype of Sur_Δ3*cpdA*. In the reciprocal mix, Sur_Δ3*cpdA* majority declined by ~7 fold whereas ancestor minority declined by ~3 fold (Fig. 3B; paired t-test comparing slopes of competing populations, N =4, P= 0.008). One order of magnitude decline of either majority population was observed in control competition between ancestor pair marked with kanamycin resistance (~9 fold) or tetracycline resistance (10 fold), and therefore not taken into account while drawing inferences.

**Figure 3:**
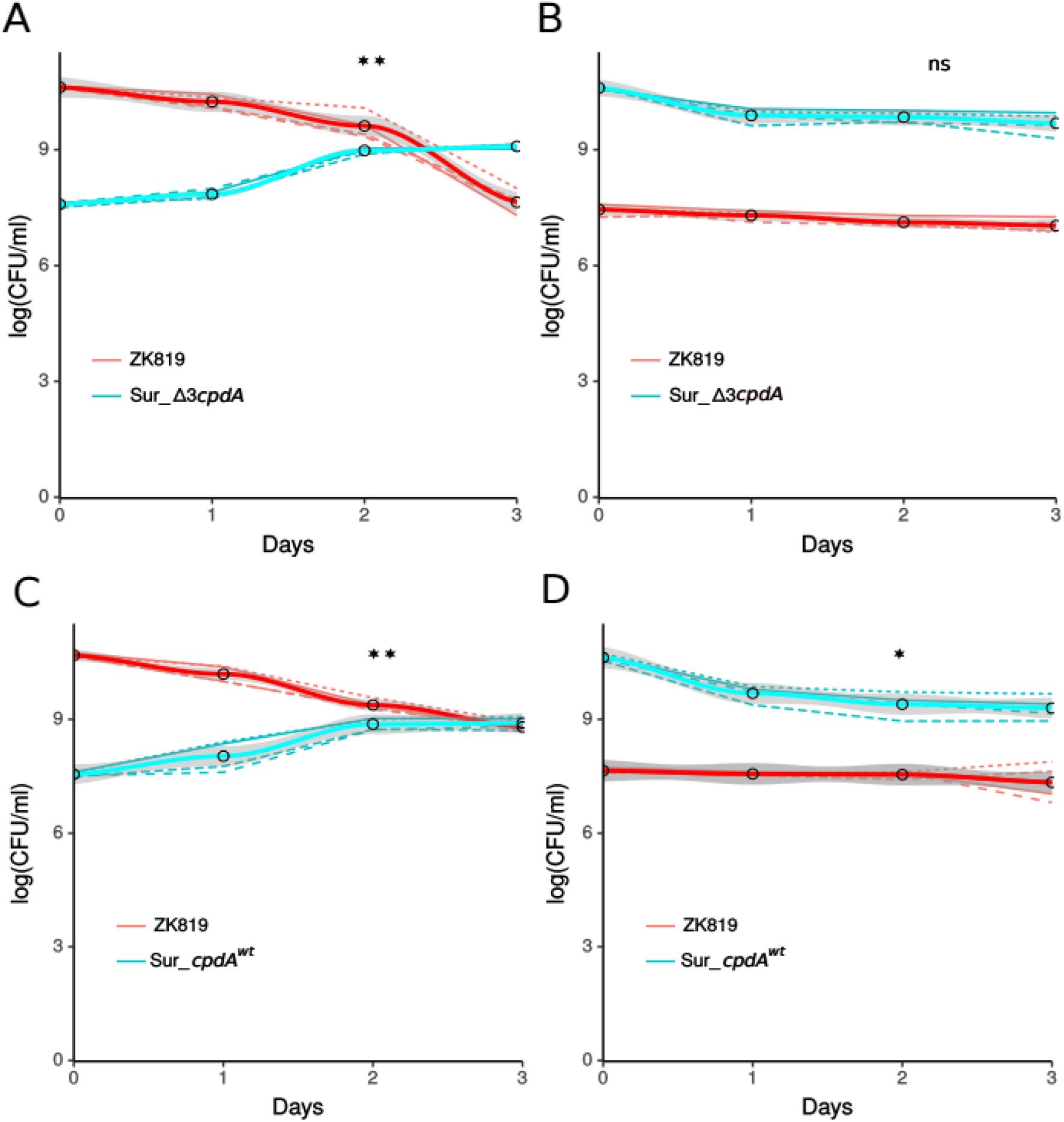
Stationary-phase competition between Sur_Δ3cpdA (blue) and ZK819 (red) (A and B).One-day-old cultures of the two strains were mixed in ratios of 1:1,000(A) and 1,000:1 (B).Competition between Sur_Δ3cpdA^wt^ (blue) and ZK819 (red) (C and D).One-day-old cultures of the two strains were mixed in ratios of 1:1,000 (C) and 1,000:1 (D).The mixed cultures were maintained in the original LB medium without the addition of fresh medium, and the growth of the two populations was monitored by scoring the number of CFU ml^−1^.**P < 0.005, *P < 0.05. The thick curve is the LOESS line fitted to the data.

To determine the contribution of Δ3*cpdA* in Sur_Δ3*cpdA* GASP, the Δ3*cpdA* allele in *Sur_Δ3cpdA* was replaced with the wild-type *cpdA* using P1 transduction. In Sur_*cpdA^wt^* versus ancestor ZK819 competition, Sur_*cpdA^wt^* minority counts increased by ~24 fold whereas ancestor majority declined by ~77 fold (Fig. 3C; paired t-test comparing slopes of competing populations, N = 4, P = 9.22 x 10^−06^). In the reciprocal mix, Sur_*cpdA^wt^* majority counts dropped by ~18 fold while the ancestor minority counts showed little change (Fig. 3D; paired t-test comparing slopes of competing populations, N = 4, P = 0.02). Thus the replacement of the Δ3*cpdA* allele by its wild type equivalent resulted in a decline in the ability of the survivor to out compete the ancestor ZK819.

### Δ3cpdA is a null allele and confers modest GASP

The cAMP phosphodiesterase encoded by *cpdA* hydrolyzes cAMP (29). In Δ3*cpdA*, a three base deletion at position 486 to 488 knocks off a leucine residue preceding a stretch of metal binding histidine residues. To test the functional status of Δ3*cpdA*, we transduced Δ3*cpdA* allele to ancestor ZK819 to create ZK819Δ3*cpdA* and measured cAMP level. For negative control, the *cpdA* gene was deleted in the ancestor ZK819 to create ZK819*cpdA*^nu^l. The cAMP level remains unaffected by the functional status of *rpoS* as indicated by the similar levels of cAMP in the presence of three *rpoS* variants namely Δ*rpoS*, *rpoS819* and *rpoS92* (Fig. 4). Deletion of *cpdA* gene in ancestor ZK819 results in ~50% increase in the level of cAMP. The natural allele Δ3*cpdA* resulted in ~40% increase in cAMP level in the transductant ZK819Δ3*cpdA* and the original survivor Sur_Δ3*cpdA,* which is comparable to that of the *cpdA* deletion strain ZK819*cpdA*^null^. Similar levels of cAMP in Sur_Δ3*cpdA* and Δ3*cpdA* transductant ZK819Δ3*cpdA* indicates that the genetic background does not influence the cAMP levels in these strains (Fig. 4).

**Figure 4:**
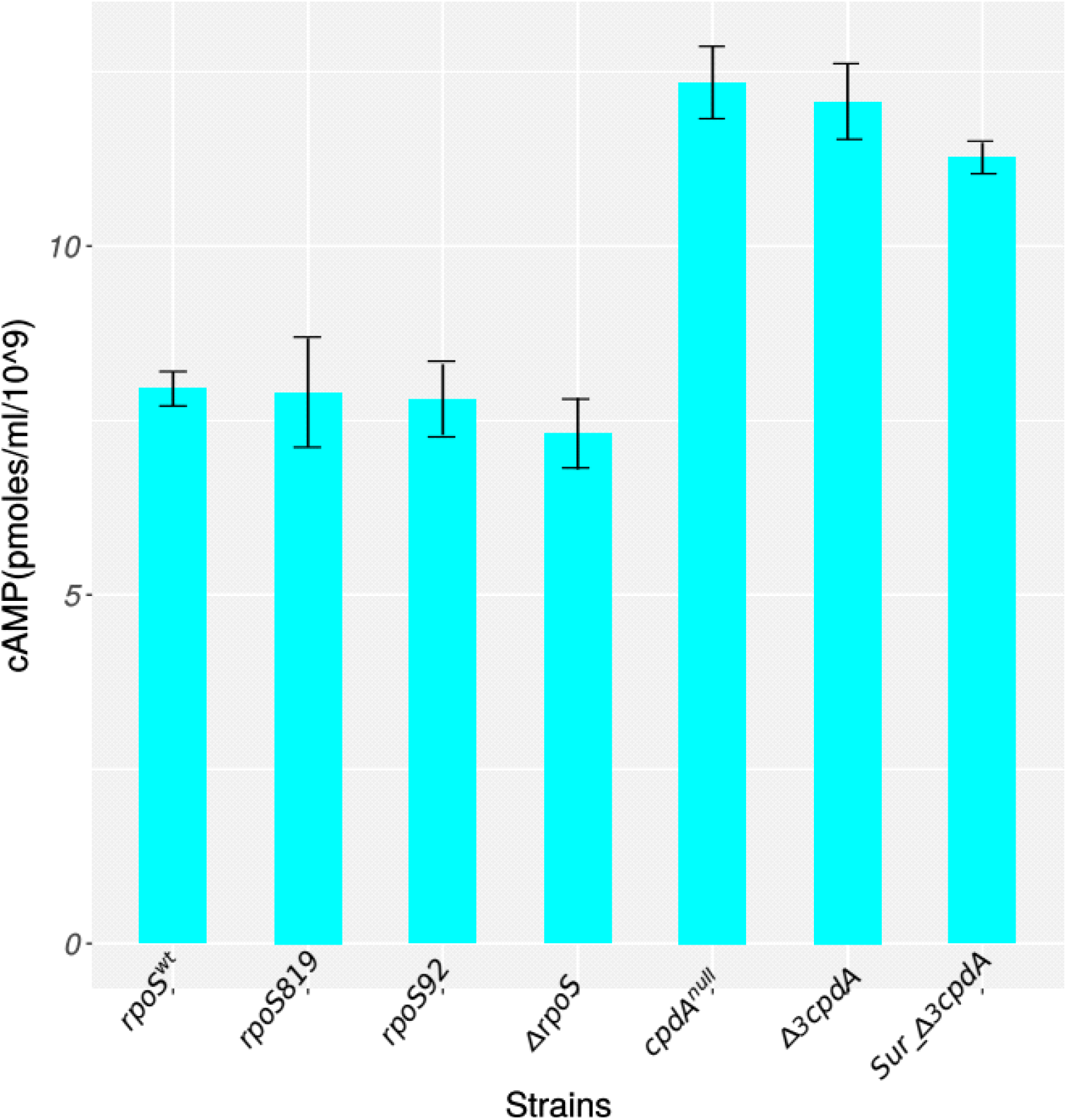
cAMP level is elevated in Δ3cpdA allele. cAMP level was measured in twelve hours old cultures with genetic background indicated using CIA cAMP kit according to manufacturer’s protocol. The error bars represent standard deviation (n = 3).

In mixed culture competitions with the ancestor ZK819, ZK819*cpdA*^null^ minority counts remained stable whereas ancestor majority declined by ~37 fold (Fig. 5A; paired t-test comparing slopes of the competing populations, N =4, P = 0.02). In the reciprocal mix, there was ~73 fold reduction in ancestor minority counts while ZK819*cpdA*^null^ majority counts dropped by ~6 fold (Fig. 5B; paired t-test comparing slopes of the competing populations, N =4, P = 0.06). In Δ3*cpdA* transductant ZK819Δ3*cpdA* versus ZK819 competition, ZK819Δ3*cpdA* minority declined by ~9 fold whereas ancestor majority counts dropped by ~186 fold (Fig. 5C; paired t-test comparing slopes of the competing populations, N =4, P = 0.002). In the reciprocal mix, ZK819Δ3*cpdA* majority counts dropped by 59 fold and ancestor minority declined by ~235 fold (Fig. 5D; paired t-test comparing slopes of the competing populations, N =4, P = 0.03). These competition assays demonstrate that loss of *cpdA* function confers modest GASP.

**Figure 5:**
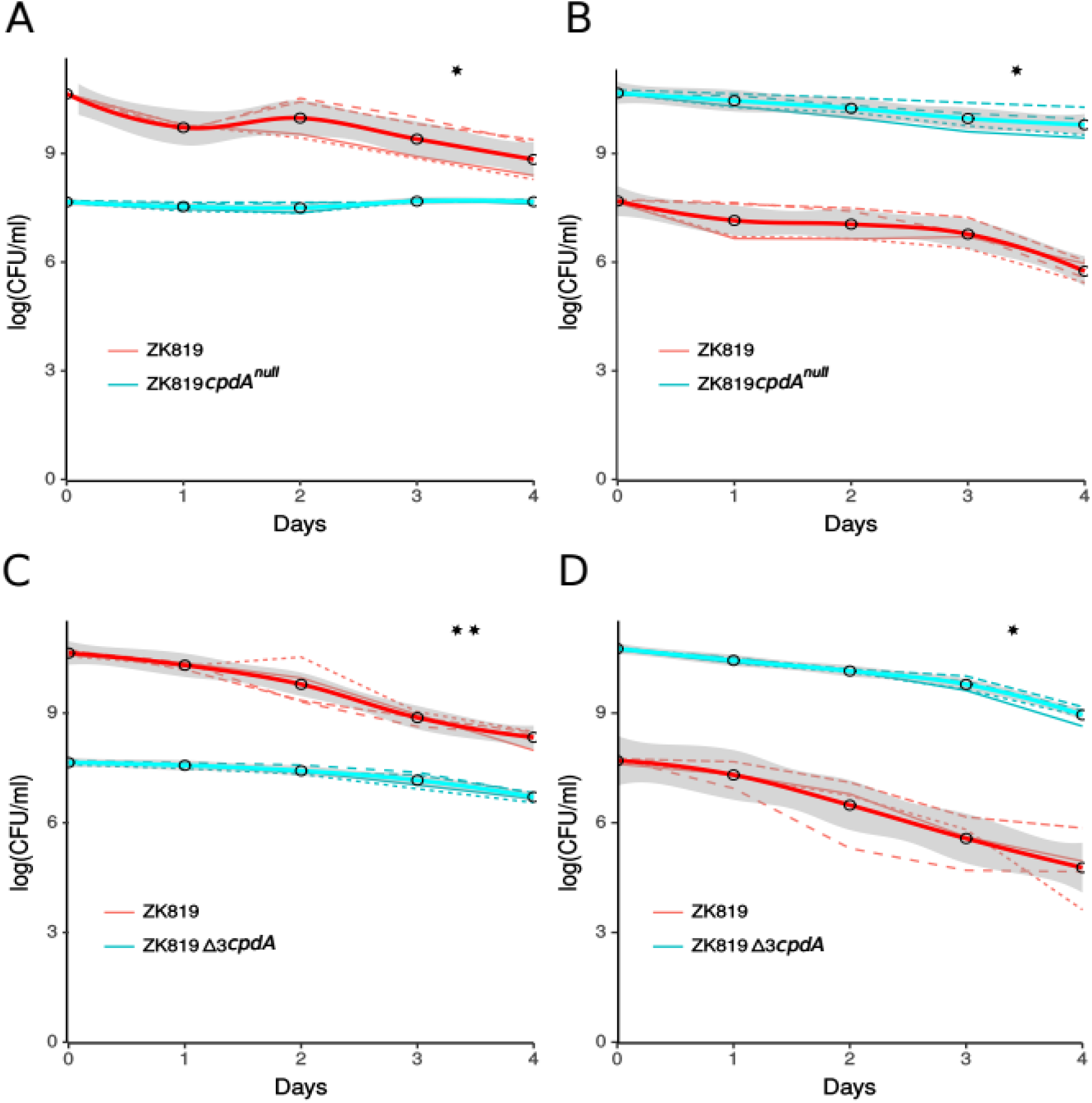
Stationary-phase competition between (A and B) ZK819 cpdAnull (blue) and ZK819 (red). One-day-old cultures of the two strains were mixed in ratios of 1:1,000(A) and 1,000:1 (B).Competition between ZK819Δ3cpdA (blue) and ZK819 (red) (C and D). One-day-old cultures of the two strains were mixed in ratios of 1:1,000 (C) and 1,000:1 (D).**P < 0.005, *P < 0.05.The mixed cultures were maintained in the original LB medium without the addition of fresh medium, and the growth of the two populations was monitored by scoring the number of CFU ml^−1^. The thick curve is the LOES line fited to the data.

### Δ3cpdA is epistatic to rpoS92 in ancestor genetic background

In previous sections, we showed that *rpoS92* transductant ZK92 displays GASP. Sur_*cpdA^wt^* partially retains GASP when Δ3*cpdA* was replaced with *cpdA^wt^*. Sur_*cpdA^wt^* genome harbours *rpoS92* along with several other mutations. To determine a potential epistatic interaction between Δ3*cpdA* and *rpoS92*, Δ3*cpdA* was transduced to ZK92 to create the double mutant ZK92Δ3*cpdA*. The double mutant ZK92Δ3*cpdA* was competed against the ancestor ZK819 in a 1:1000 reciprocal mix. During competition, the double mutant ZK92Δ3*cpdA* minority counts hardly increased while the ancestor ZK819 majority declined by 21.77 fold (Fig. 6A; paired t-test comparing slopes of the competing populations, N = 4, P = 0.02). In the reciprocal experiment, ZK92Δ3*cpdA* majority declined by ~4 fold whereas ancestor minority declined by ~17 fold (Fig. 6B; paired t-test comparing slopes of the competing populations, N = 4, P = 0.1). The presence of Δ3*cpdA* in ZK92 masks the *rpoS92* mediated GASP of ZK92 indicating that Δ3*cpdA* is epistatic to *rpoS92* in ancestor background (compare Fig. 1 with Fig. 6). The absence of this epistatic interaction in the original survivor *Sur_Δ3cpdA* suggests the role of the additional mutation/s present on its genome in neutralizing this epistatic interaction.

**Figure 6:**
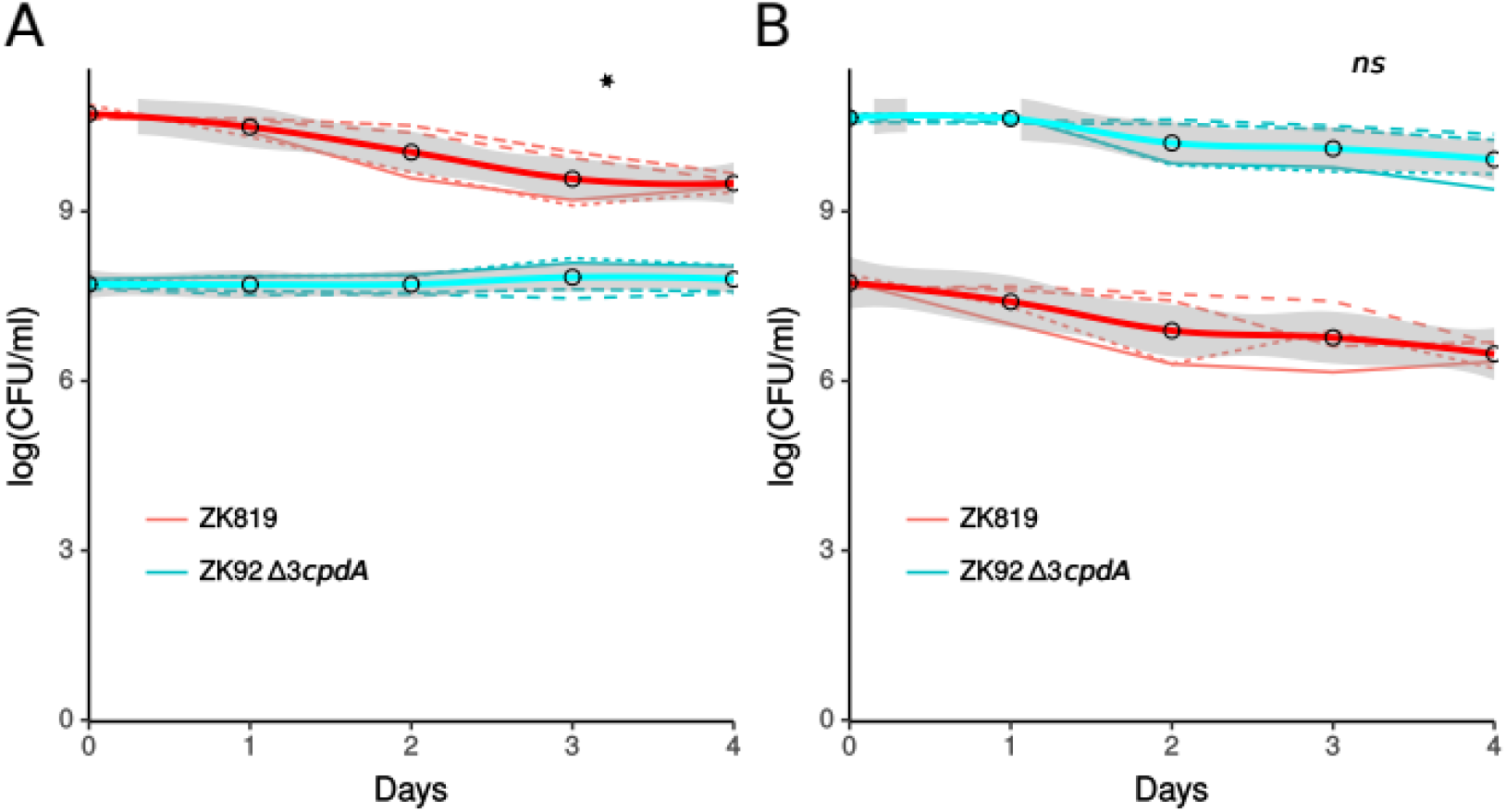
Stationary-phase competition between ZK92Δ3cpdA (blue) and ZK819 (red) (A and B). One-day-old cultures of the two strains were mixed in ratios of 1:1,000 (A) and 1,000:1 (B). The mixed cultures were maintained in the original LB medium without the addition of fresh medium, and the growth of the two populations was monitored by scoring the number of CFU ml^−1^. **P < 0.005, *P < 0.05, ns > 0.05; not significant. The thick curve is the LOESS line fited to the data.

### Age of spent medium influences rpoS92 and cpdA fitness

The competitions discussed in the above sections typically represent the environment prevailing during the first week of our evolution experiment. While the competition assays performed in the stationary-phase reflect the correct environmental context for the emergence of *rpoS92*, most of the *cpdA* mutations were observed during the second and third weeks of the original evolution experiment. A reasonable approach to recreate a later stage approximation of environmental context is to generate spent medium by growing the ancestral culture till the desired age and use this medium for competition experiments. We chose ten day and twenty one day old conditioned medium for these experiments. In spent medium competition, the *rpoS92* transductant ZK92 minority declined by ~4 fold and the ancestor ZK819 majority declined by ~56 fold (Fig. 7A; paired t-test comparing slopes of the competing populations, N = 4, P = 0.02). In the reciprocal mix, ZK92 majority declined by ~64 fold whereas ancestor ZK819 minority counts dropped by ~1461 fold (Fig. 7B; paired t-test comparing slopes of competing population, N = 4, P = 0.004). Thus, in ten day old spent medium, the competitive fitness of ZK92 is still higher than that of the ancestor, although the degree to which ZK92 in minority closes the gap with ZK819 majority is less in the spent medium than in fresh LB.

**Figure 7:**
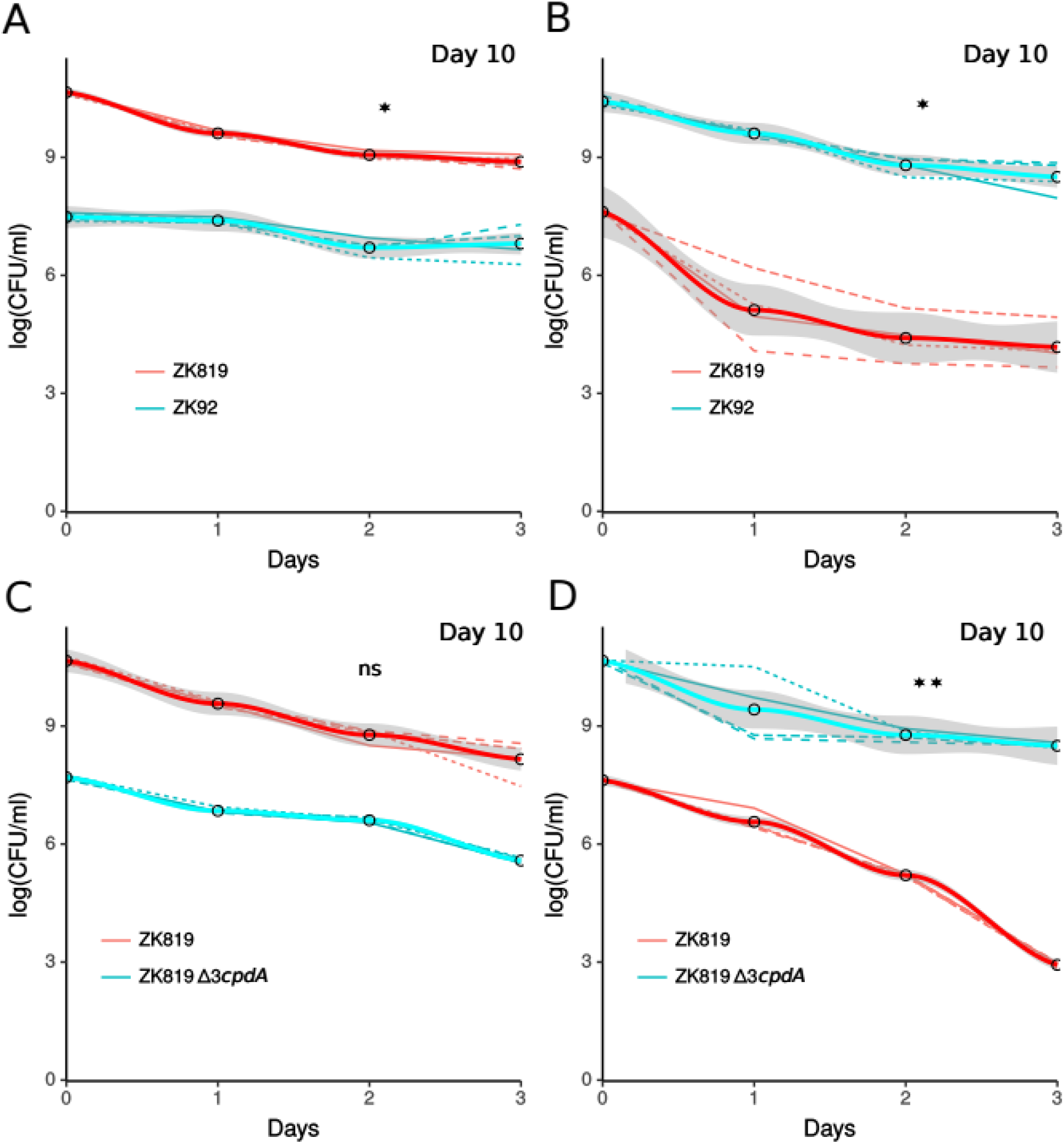
Competition between ZK92 (blue) and ZK819 (red)(A and B).One-day-old cultures of the two strains were mixed in ratios of 1:1,000 (A) and 1,000:1 (B) in spent medium derived from a 10 Days old parent culture. Competition between ZK819Δ3cpdA (-) and ZK819 (-) (C and D).One-day-old cultures of the two strains were mixed in ratios of 1:1,000 (C) and 1,000:1 (D) in spent medium derived from a 10 Days old parent. **P < 0.005, *P < 0.05, not significant (ns)> 0.05. The thick curve is the LOESS line fitted to the data.

In Δ*3cpdA* transductant ZK819Δ3*cpdA* versus ancestor ZK819 competition in ten day old spent medium, ZK819Δ3*cpdA* minority counts declined by ~130 fold and the ancestor ZK819 majority declined by ~228 fold (Fig. 7C; paired t-test comparing slopes of the competing populations, N =4, P = 0.1). In the reciprocal mix, ZK819Δ3*cpdA* majority declined by ~144 fold whereas ancestor ZK819 minority declined by 49110 fold (Fig. 7D; paired t-test comparing slopes of the competing populations, N = 4, P = 0.0002). In ten day old spent medium, there is a reduction in ZK819Δ3*cpdA* and ancestor ZK819 fitness as reflected by the decline of the competing populations both as majority or minority fraction. However, ancestor fitness is severely reduced when present in minority with more than four orders of magnitude decline as opposed to nearly two and half orders of decline of ZK819Δ3*cpdA* minority.

During the double mutant ZK92Δ3*cpdA* versus ancestor ZK819 competition in ten day old spent medium, ZK92Δ3*cpdA* minority counts remained stable while ancestor ZK819 majority counts dropped by ~493 fold (Fig. 8A; paired t-test comparing the slopes between competing populations, N = 4, P = 0.001). In the reciprocal mix, ZK92Δ3*cpdA* majority declined by ~29 fold and the ancestor minority declined by 653818 fold (Fig. 8B; paired t-test comparing the slopes between competing populations, N = 4, P = 0.001). The ten day old spent medium competition of ZK92Δ3*cpd* indicates that the competitive fitness of the double mutant ZK92Δ3*cpd* competitive fitness against ancestor is higher compared to that of the either of the single mutants ZK92 or ZK819Δ3*cpdA*.

**Figure 8:**
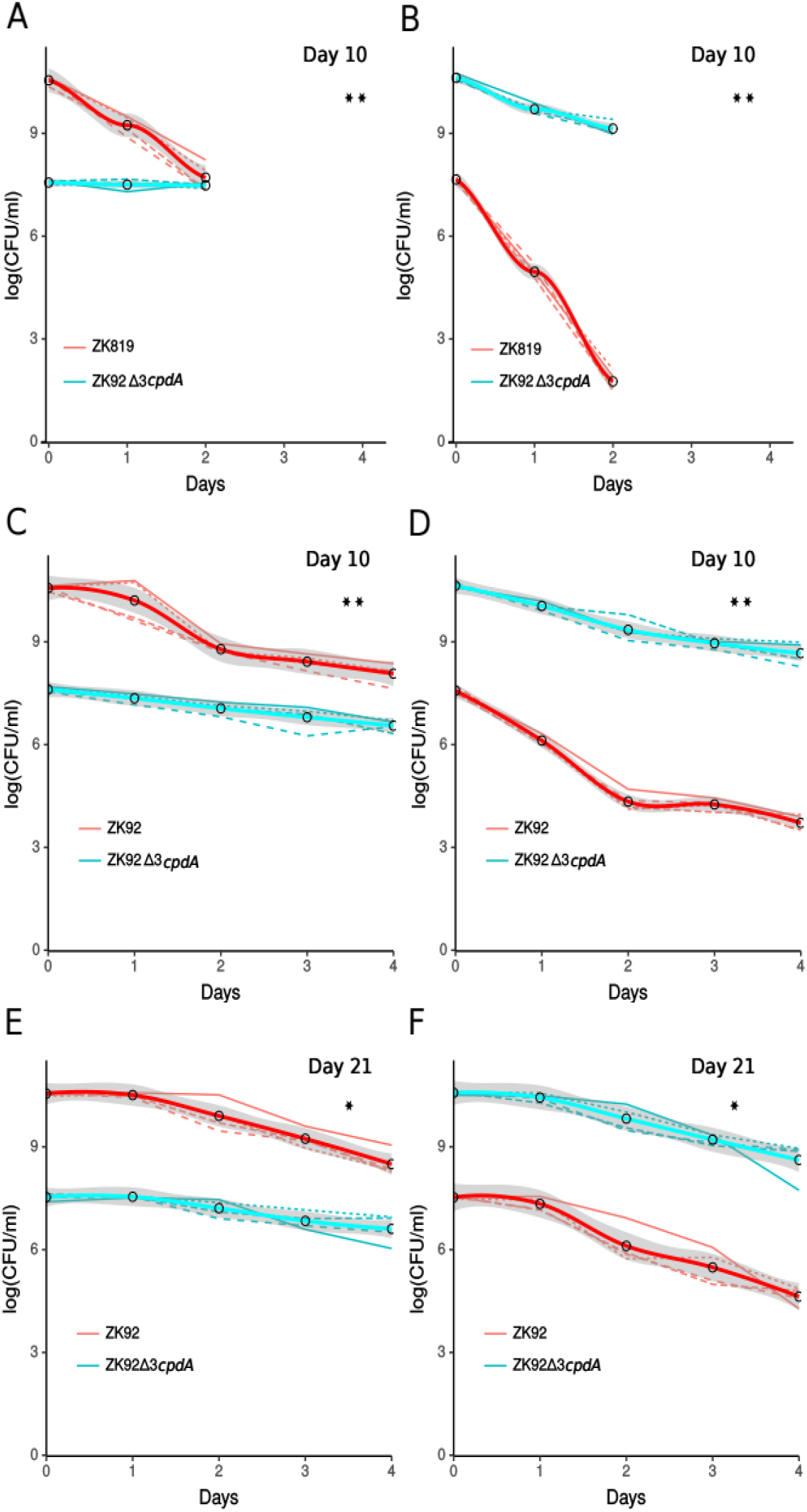
Competition between ZK92Δ3cpdA (blue) and ZK819 (red) (A and B). One-day-old cultures of the two strains were mixed in ratios of 1:1,000 (A) and 1,000:1 (B) in spent medium derived from a 10 Days old parent culture. Competition between ZK92Δ3cpdA (blue) and ZK92 (red) (C and D).One-day-old cultures of the two strains were mixed in ratiosof1:1,000(C) and 1,000:1 (D) in spent medium derived from a 10 Days old ZK819 culture. Competition between ZK92Δ3cpdA (blue) and ZK92 (red) (E and F).One-day-old cultures of the two strains were mixed in ratios of 1:1,00 (E) and 1,00:1 (F) in spent medium derived from a 21 days old ZK819 culture.**P < 0.005, *P < 0.05, not significant (ns) > 0.05. The thick curve is the LOES line fitted to the data.

We tested the fitness of the double mutant ZK92Δ3*cpdA* against its more plausible competitor ZK92 in ten day and twenty one day old spent medium. In ten day old spent medium competition between ZK92Δ3*cpdA* and ZK92, ZK92Δ3*cpdA* minority counts declined by ~10 fold and ZK92 minority counts declined by ~270 fold (Fig. 8C; paired t-test comparing slopes of the competing populations, N =4, P = 0.002). In the reciprocal mix, ZK92Δ3*cpdA* majority declined by ~77 fold and ZK92 minority declined by ~6567 fold (Fig. 8D; paired t-test comparing slopes of the competing populations, N =4, P = 0.001). In twenty-one day old spent medium competition, ZK92Δ3*cpdA* minority counts declined by ~6 fold whereas the ZK92 majority declined by ~77 fold (Fig. 8E; paired t-test comparing slopes of the competing populations, N = 4, P = 0.04). In the reciprocal mix, ZK92Δ3*cpdA* majority declined by ~60 fold and ZK92 minority declined by ~747 fold (Fig. 8F; paired t-test comparing slopes of the competing populations, N = 4, P = 0.01). The competitive fitness of the double mutant ZK92Δ3*cpdA* is higher than ZK92 in both ten day as well as twenty-one day old spent medium. However, ZK92 fitness improved in twenty-one day old spent medium with more than ten fold reduction in the population decline. Spent medium competition experiment trends show that Δ3*cpdA* arrests the decline in fitness of rpoS92 in deep stationary-phase.

## Discussion

Laboratory evolution experiments combined with genome sequencing has made it possible to catalog mutations and understand adaptation. The growth advantage in stationary phase (GASP) paradigm, first demonstrated in *E. coli,* illuminates a common survival strategy under persistent starvation (10, 12, 13, 14). The GASP mutants establish continuous growth and death cycles under growth limiting environment of prolonged stationary-phase (10, 48). The dynamics of competitive fitness of GASP mutants is reflected in the different GASP flavours displayed by the competing populations (48). One of the earliest GASP alleles to be identified from the survivors of prolonged stationary-phase was a new allele of stationary-phase sigma factor named *rpoS819* (10).

In an earlier study, we had reported the mutation spectrum obtained during a second round of long-term stationary phase evolution of ZK819, an *E. coli* strain with *rpoS819* (39). A key observation of this study, as well as other recent studies of evolution in prolonged stationary phase in complex medium is the prevalence of parallel evolution with mutations in genes that have pleiotropic effects (39, 50-51). For example, in a very similar prolonged stationary phase experimental evolution setup like ours, though of longer duration, Hershberg and colleagues had reported multiple mutations in the RNA polymerase core subunits, but not in *rpoS*, which is usually one of the first gene to be hit by mutations under nutrient limitation (51). In addition to parallel mutations at different RNA polymerase subunits, we had observed parallel evolution at stationary-phase sigma factor *rpoS* and cAMP-phosphodiestersae *cpdA* in our one month old long-term evolution (39). In the present study, we characterized new alleles of *rpoS* and *cpdA* namely *rpoS92* and Δ3*cpdA* and investigated their fitness potential during prolonged stationary-phase.

We show that RpoS status fluctuates at a genetic level through 30 days of stationary phase, and that additional mutations in the cAMP phosphodiesterase CpdA modulates the fitness of RpoS variants. Heterogeneity at *rpoS* has been frequently observed particularly in several laboratory strains of *E. coli*. Modulation of *rpoS* activity has been shown to facilitate a trade-off between general stress response and nutritional competence (6-8, 21, 23). The *rpoS819* mediated GASP has been directly correlated with enhanced ability to scavenge amino acids, the key nutrients released by dead cells. This can be modulated by altering environmental factors such as pH of the medium (11, 15, 52). Strong selection for *rpoS92,* which partially recovers RpoS activity, indicates further tuning of RpoS activity. As stationary-phase deepens in a closed system such as batch cultures, nutrition may drop below a threshold to support growth. Bringing gene-expression closer to that of the wild type RpoS, RpoS92 might increase the general stress response while compromising the nutritional competence. Indeed in spent medium, though *rpoS92* failed to manifest GASP, it showed better survival advantage against the *rpoS819* ancestor.

Mutations that abrogated the activity of CpdA appeared in multiple lines during the second half of our GASP experiment. The CpdA mutation (Δ3*cpdA*) studied here appeared to enhance the survival of *rpoS92* in spent medium competitions, whereas it dampened the competitive fitness of *rpoS92* against *rpoS819* in fresh medium competitions. Therefore, the selective advantage conferred by Δ3*cpdA* appears to take effect only late in stationary phase. However our observation that the negative effect of Δ3*cpdA* on *rpoS92* is not prominent in the genetic background of the original survivor that carried additional mutations beyond *rpoS92* and Δ3*cpdA*, suggests that these mutations may also play a role in stationary phase growth / survival.

To gain a molecular mechanistic insight underlying adaptaion, we performed RNAseq on *cpdA* strains, which however failed to identify any significant differentially expressed genes when compared to an otherwise isogenic background. Thus, we did not find evidence for increased nutritional competence via cAMP-CRP mediated regulation (unpublished data). However, these experiments were performed in fresh LB, and whether a different set of results will be obtained in a consistent manner in spent medium RNA-seq experiments remains an open question. There is increasing evidence supporting a role for cAMP in negative regulation of persistence, a phenotype observed under extreme stress (53-55). Identifying the specific background mutation/s modulating the epistatic interaction between Δ3*cpdA* and *rpoS92* will provide additional insights to the genetic regulation of proliferation versus non-proliferation mechanisms under severe stress.

## Accession Numbers

Genome sequence data for Sur_Δ3*cpdA* is available from the Sequence Read Archive (SRA) database under the accession no. SRP094816 (https://www.ncbi.nlm.nih.gov/search/?term=SRP094816). RNA sequencing data from this study are available from the Gene Expression Omnibus (GEO) database under the accession number GSE119046 (https://www.ncbi.nlm.nih.gov/gds/?term=GSE119046).

## Acknowledgment

We thank S. Mahadevan for discussions. SC was supported by DST-SERB young scientist grant SB/YS/LS-148/2014. ASNS was supported by DBT grant BT/PR3695/BRB/10/979/2011, Ramanujan fellowship SR/S2/RJN-49/2010 to A.S.N.S. and a DBT Wellcome Trust India Alliance Intermediate fellowship IA/I/16/2/502711.

---

Supplementary Table 1: Additional mutations in original survivor Sur_Δ3*cpdA*

